# Single and population coding of taste in the gustatory-cortex of awake mice

**DOI:** 10.1101/575522

**Authors:** David Levitan, Jian-You Lin, Joseph Wachutka, Narendra Mukherjee, Sacha B. Nelson, Donald B. Katz

## Abstract

Electrophysiological analysis has reveals much about the broad coding and neural ensemble dynamics that characterize gustatory cortical (GC) taste processing in awake rats, and about how these dynamics relate to behavior. With regard to mice, meanwhile, data concerning cortical taste coding have largely been restricted to imaging—a technique that reveals average levels of neural responsiveness, but that (currently) lacks the temporal sensitivity necessary for evaluation of fast response dynamics; furthermore, the few extant studies have thus far failed to provide consensus on basic features of coding. We have recorded the spiking activity of ensembles of GC neurons while presenting representatives of the basic taste modalities (sweet, salty, sour and bitter) to awake mice. Our first central result is the identification of similarities between rat and mouse taste processing: most mouse GC neurons (~66%) responded distinctly to multiple (3-4) tastes; temporal coding analyses further reveal, for the first time, that single mouse GC neurons sequentially code taste identity and palatability—the latter responses emerging ~0.5s after the former—with whole GC ensembles transitioning suddenly and coherently from coding taste identity to coding taste palatability. The second finding is that spatial location plays very little role in any aspect of taste responses—neither between- (anterior-posterior) nor within-mouse (dorsal-ventral) mapping revealed anatomical regions with narrow or temporally simple taste responses. These data confirm recent results showing that mouse cortical taste responses are not “gustatopic,” but also go beyond these imaging results to show that mice process tastes through time.

## Introduction

Mice are the most commonly used subjects in vertebrate neuroscience research. The use of mice affords researchers relatively easy access to the molecular and genetic underpinnings of network activity, learning-related plasticity, and behavior (Stevens, 1996; Kandel et al., 2014). As taste is a particularly good sensory system with which to study these three topics (see below and Carleton et al., 2010; Maffei et al., 2012), it is surprising that there has been almost no extensive analysis of central taste electrophysiology in awake mice (although see Kusamoto-Yoshida, et al., 2015). There have been several *in vivo* analyses of mouse gustatory cortical (GC) taste responses based on calcium imaging data, but the results of these analyses, while exciting, have been difficult to reconcile with one another—work from one group suggests extreme spatial separation of responses to different tastes (Chen et al., 2011; Peng et al., 2015), while work from other groups suggest broad and non-mapped responses (Fletcher et al., 2017; Livneh et al., 2017; Lavi et al., 2018).

Furthermore, calcium imaging in general lacks the temporal sensitivity necessary for evaluation of fast single-trial response dynamics, a potentially important limitation given the evidence that GC taste responses in awake rats contain precisely such dynamics. More specifically, the rat work has reliably revealed that: 1) responses to multiple tastes appear not only in the same region but also within the same single neurons (Katz et al., 2001; Fontanini & Katz, 2006; Jezzini et al., 2013; Accolla et al., 2007; Samuelsen et al., 2013; Samuelsen & Fontanini, 2017); 2) these responses progress through a series of firing-rate “epochs,” coding in turn the presence, physical properties, and palatability of a taste across 1-1.5s (Katz et al., 2001; Bahar et al., 2004; Sadacca et al., 2012; Maier & Katz, 2013; Samuelsen et al., 2013); 3) late-epoch palatability-related firing is uniquely affected by experience-driven shifts of taste palatability (Bahar et al., 2004; Fontanini & Katz, 2006; Grossman et al., 2008; Moran and Katz., 2014); and 4) in single trials the onset of palatability is a sudden, coherent network phenomenon, the timing of which predicts and likely drives behavior (Jones et al., 2007; Sadacca et al., 2016; Li et al., 2016). A test of whether mouse GC neurons respond in a like manner has not yet been performed, and cannot be performed using imaging data.

For the current study, we have recorded the responses of small ensembles of GC neurons in awake mice to presentations of stimuli representing four major taste modalities (sweet, salty, sour and bitter). Our data reveal, first of all, that taste responses in mouse GC recapitulate the dynamics of taste responses in rat GC: taste delivery evoked broad and dynamic responses that coded first taste identity and then taste palatability across 1.0-1.5 seconds; furthermore, these single-neuron dynamics reflected coherent ensemble transitions—in individual trials, multiple neurons in each ensemble hopped from one firing rate to another coherently, mid-trial. We took care to map the spatial extent of GC, and were therefore also able to show that these response properties are observed uniformly along the anterior-posterior and dorsal-ventral axes; there were no regions of noticeably sparse responsiveness, and no regions in which responses failed to reflect the general dynamics.

These results offer novel insight into mouse cortical taste processing: they support recent imaging studies suggesting that mouse taste responses are broad and non-gustatopic (Fletcher et al., 2017; Livneh et al., 2017; Lavi et al., 2018), but they also go beyond this to suggest that mouse cortical taste processing is dynamic, and that therefore “breadth” measures fail to fully describe GC taste function in awake mice (i.e., breadth meaningfully changes across the “stages” of the response). These insights can be used, in conjunction with tools available only in the mouse (among mammals), to delve into the undoubtedly rich relationship between molecular and network analyses of sensory function.

## Material and Methods

### Subjects

The experimental subjects were C57BL/6J (n = 2) and Stk11tm1.1Sjm (StK11; n = 4) mice. The mice were purchased from Jackson Laboratories (Bar Harbor, ME). Stk11f/f mice have conditional floxed allele between exon 3 and exon 6, which was inactive through the entire set of experiments. These mice were backcrossed after development by the donating investigator for at least 4 generations to C57BL/6J mice and were bred to C57BL/6J for at least one generation upon arrival to Jackson Laboratories (see also Nakada et al., 2010). After the mice were purchased from Jackson Laboratories they were further backcrossed for 3-4 generation to C57BL/6J before establishing a colony. Upon arrival, the mice were placed on a 12-hour light-dark cycle, and given *ad libitum* access to food and water except during experimentation, at which time water access was restricted (see “Fluid delivery protocol”) while food remained available *ad libitum* (note that animals reliably consume less food when thirsty).

No behavioral or neural taste response differences distinguished the strains; they are therefore collapsed into a single group for purposes of our experiments. Experiment procedures started once mice were 60-80 days old. All procedures were approved by the Brandeis University Institutional Animal Care and Use Committee (IACUC) and in accord National Institutes of Health guidelines.

### Surgery

Mice were anesthetized with an intraperitoneal (ip) injection of ketamine/xylazine mix (KX; 20mg/ml ketamine, 2.5 mg/ml Xylazine, and 0.5 mg/ml Aceptomazine; total injection volume 5μl/g), and stabilized in a stereotaxic frame. A midline incision on the scalp exposed the skull, and 2 trephine holes were made dorsal to GC. Multi-electrode bundles (16 formvar-coated, 25-μm in diameter nichrome wires) were gradually lowered into the holes, to a spot just above GC: the AP position was ~1.2 mm in front of bregma for anterior GC and ~0.4 mm for posterior GC; the ML positioning was ±3 mm for all mice. Bundles were then lowered to a DV of −2.25 mm from the *pia mater* (prior to recording sessions, bundles were further lowered by 0.25-0.5 mm to reach dorsal GC, and 0.75-1.00 mm to reach ventral GC; see Figure 1 for examples of the identification of the various anatomical locations) and cemented to the skull, along with one intraoral cannula (IOC; flexible plastic tubing inserted into the cheek to allow controlled delivery of tastes to the tongue of awake mice, see below).

**Figure 1.**
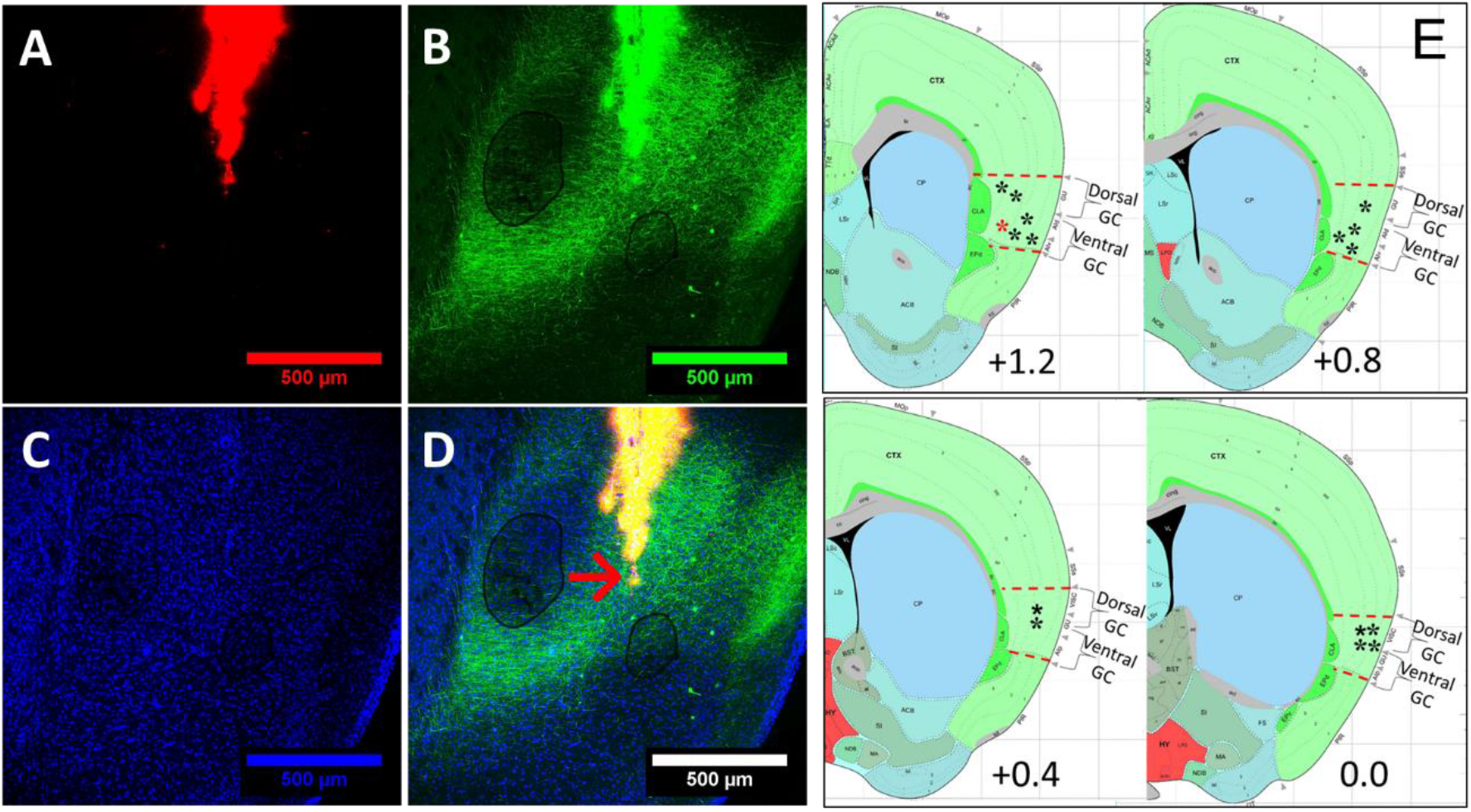
Localization of electrode bundles in mouse GC. **A**. To determine the recording site in GC, electrode bundles were coated with Di-l (a red fluorescent tracer) which later showed the track traveled by the multielectrode array in GC. **B**. To assist the separation between dorsal and ventral subregions of GC, BLA was infected with AAV2/5::GFP; resultant GFP-labeled axons were then visible in ventral GC. **C** Background staining of the same slice with DAPI (a nonspecific nuclear dye). **D**. In a triple-labeled slice, an arrow indicates the final location of the tips of the electrodes. **E**. A schematic representation of all electrode bundles (asterisks; red asterisk represent electrode from D) on top of a schematic of a coronal slice containing GC (Bregma +1.6 mm to Bregma 0.0 mm; every 0.4 mm were collapsed to a single representative image; Allen Institute for Brain Science, mouse brain atlas, P56, coronal). The red asterisk specifically corresponds to the electrode from D.

In a subset of mice, basolateral amygdala (BLA; AP= −1.4, ML= ±3.4, 200 nl injections at DV= 4.3 mm and 4.6 mm from the dura) was infected with AAV2/5::Camk2α-GFP to visualize BLA axons in the ventral GC. This was done to make the separation of dorsal from ventral sub-regions of GC easier to visualize.

Mice were given 7 days to recover from the surgery. During the first 3 days of this recovery, injections of meloxicam (2 mg/kg), ip) and penicillin (0.1 ml), both administered subcutaneously, provided pain and infection management, respectively.

### Fluid delivery protocol

To accommodate the requirement of electrophysiology recording, experiments were conducted in in a Faraday cage constructed with standard aluminum insect netting with dimensions as 26 cm × 24 cm × 33 cm (length x width x height). Fluid was delivered through a nitrogen-pressurized system of polyethylene tubes; flow was controlled by solenoid valves opened by a Raspberry Pi computer installed with customable python scripts that control the timing and quantity of each taste delivery (construction details and code available on request from [https://github.com/narendramukherjee/blech_clust]. During the experiment, mice were allowed to move about the chamber without restraint.

For three days following recovery from surgery, sessions consisted of 60 deliveries of 15- μl aliquots (hereafter, “trials”) of water across 30 min. This procedure habituated the mice to the experimental environment, and to IOC delivery of fluid. Following the last water-delivery session, electrodes were lowered 250-1000 μm and mice were returned to home cage.

On the following day, the recording experiment commenced. The procedure was identical to that described above, except that water was replaced with 4 different taste stimuli, which were sweet (0.2 M sucrose), salty (0.1 M sodium chloride), sour (0.02 M citric acid), and bitter (0.001 M quinine). A total of 15 trials of each taste were delivered, in random order (sampling without replacement). These tastes and concentrations were chosen to provide compatibility to our prior rat research, and because they provided a broad range of both quality and palatability.

A subset of mice (N=4) received two identical recording sessions separated by one day. In these cases, the electrodes were lowered ~250-500 μm immediately after the first recording session. When data from these two sessions were compared (as a specific part of the study, see below), no substantive between-day differences were noted; the data from the two sessions were therefore otherwise combined.

### Acquisition of electrophysiological data

Voltage signals from the micro-electrodes were sampled at 30 kHz, digitally amplified and filtered, and saved to the hard drive of an Ubuntu computer connected to an Intan recording system (RHD2000 Evaluation System and Amplifier Boards; Intan Technologies, LLC, LA). We retained all waveforms from these raw voltage signals, and sorted the waveforms into distinct units using a semi-supervised spike sorting strategy: recorded voltage data were filtered between 300-3000Hz, grouped into potential clusters by a Gaussian Mixture Model (GMM) and refined manually (to enhance conservatism) by the experimenters (for details see Mukherjee et al., 2017). A total of 185 single units were isolated for the experiments across 10 sessions. As has been done many times previously, isolated units were categorized as being either putative pyramidal neurons or putative interneurons on the basis of wave shape and average inter-spike interval (e.g., Bartho et al., 2004; Sirota et al., 2008).

### Statistical analysis of neural data

*“Taste responsiveness”*. A single neuron was deemed “taste-responsive” if firing rates changed with taste delivery. Paired t-tests were conducted to determine whether the evoked firing rate (2 seconds post-delivery) significantly differed from its baseline firing rate (2 seconds prior taste delivery). Alpha level was set at *p* < 0.05. This analysis was first performed on firing averaged across taste (providing a general quantification of the neuron’s sensitivity to taste input) and then again for each individual taste response.

Of course, this analysis (like all of those described below, and all of those used in every electrophysiology study) is imperfect. For one thing, this particular analysis assumes independence of samples, which strictly speaking trials are not. Fortunately, our test is robust to this assumption, and any lack of trial-to-trial independence is likely to make responses less taste-distinctive; furthermore, the tests below, which suffer even less from this assumption, provide a converging measure of the validity of our results. Similarly, while this analysis generates a relatively large number of statistical comparisons (a ubiquitous issue for studies of simple coding), the combination of this with the more unified, more conservative analyses described below ensures valid recognition of the nature of mouse GC taste coding.

*“Taste specificity”*. The above analysis provides one estimate of the number of tastes to which a neuron responds. This fails to provide a rigorous description of the taste-specificity of the neuron’s firing, however—it is now well-recognized that the reaching of conclusions regarding differences between responses using only separate tests of each response (e.g., concluding that “because the neuron responds to both X and Y, its responses to X and Y are the same” or that “because the neuron responds significantly to X but not to Y, its responses to X and Y are significantly different”) is statistically invalid (Nieuwenhuis, et al., 2011).

We instead tested (as we have using rat data, Li et al., 2013; Piette et al., 2012; Sadacca et al., 2012) whether a single-neuron’s responses were “taste specific” *via* direct comparison of the firing rates elicited by the 4 tastes. Specifically, taste-induced firing was divided into four consecutive 500-ms bins, and a repeated-measure ANOVA was conducted with taste and time as within-subject variables; a neuron’s responses were categorized as taste-specific if either the main effect of taste or the taste X time interaction was significant (p < 0.01): the former significant effect indicates simple taste specificity— at least one taste induced an amount of spiking that differed significantly from that driven by at least one other taste; the latter indicates “temporal coding” (in quotes because the suggestion is not that taste identification comes from analysis of the time-course of firing, but that cortical taste processing is poorly characterized using only overall firing rate averages, see Discussion)—that the time course of the response to one taste was different from the time course of the response to at least one other taste (note that a set of responses could be identical in average rate but differ in time course, in which case the taste main effect would be non-significant while the taste x time interaction would be significant). Post-hoc tests (Tukey’s HSD) revealed precisely which responses differed from which.

Note that this analysis, while non-standard in the field (largely because our approach is unusual in collecting multiple replicates of data with high temporal sensitivity), avoids the above-described pitfalls to which other analyses of response breadth fall prey, allowing uncommonly valid statements of how single-neuron responses to multiple tastes differ from one another. It is beyond the scope of this report to catalog and critique techniques used to assess neural coding in the taste field, but suffice it to say that all have their limitations, and that the one described here provides a direct assessment of a subtle form of the question “is the firing of this neuron taste-specific” without introducing specific biases toward answering that question with a “yes.”

*“Entropy (H)”*. We employed the oft-used entropy (H) metric as a direct evaluation of the breadth of single-neuron tuning. H was computed as previously described by Smith and Travers (1979) using the following formula (see also Wilson and Lemon, 2013):

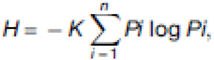

where K is a constant (K = 1.66 for a 4-taste battery), and P is the proportional response to each taste (i). A low H value indicates higher selectivity (H=0 means that a unit responds purely to one taste), whereas a high H value represents broader tuning (H=1 means that a unit responds to all tastes). Taste evoked responses used for entropy analyses were calculated by subtracting mean firing rates of pre-taste delivery (2 seconds) from post-taste taste delivery (2 seconds).

Note that this entropy analysis assumes that responses are positive-going from a zero baseline, whereas neural sensory responses may be negative going as measured from a spontaneous firing baseline—that is, responses may be inhibitory. We dealt with this issue simply, by taking the absolute value of any inhibitory response for use in the calculation of an H index (Wilson and Lemon, 2013). In practice, this solution was a conservative one: inhibitory responses, which are necessarily magnitude-limited by the zero-firing floor effect, will tend to look small in this analysis, despite the fact that they reflect particularly entropic responses (those that vary from substantially below to above zero); thus the use of absolute values will most likely lead to under-estimation of a neuron’s breadth of responsiveness.

But when combined with the above analyses, entropy provides a rich description of GC neuron taste-responsiveness. For instance, a neuron that responds to all tastes equally would have high entropy and broad taste responsiveness, but would likely be responding not to taste quality *per se* but to some other aspect of the stimulus. Accordingly, its firing would almost certainly fail to be deemed taste-specific according to the taste main effect in a 2-way ANOVA.

Finally, it is worth noting that any particular H between 0 and 1 can conceivably be described to represent “broad” or “narrow” coding, depending upon one’s perspective. To enhance interpretability of H distributions, we therefore compared our results to those of identical analyses performed on simulated datasets (created based on the real data) made up of units with known breadth.

“*Time course of palatability-related responding*.” Once more taking a cue from our work on rats (e.g., Fontanini et al., 2009; Piette et al., 2012; Sadacca et al., 2012; Li et al., 2016; Sadacca et al., 2016), we used a moving-window analysis to identify time periods in which GC taste responses reflected the hedonic value of the tastes. We first determined the palatability of each of the taste used in this study for the strain of mice used here: a set of 32 mice of the same strain but separate from those used in the electrophysiology experiment were mildly water restricted for 3 days (two 30-min periods of access to water per day: one in the morning, and another in the afternoon). Across the following 4 days, mice were exposed to each of the tastes (sucrose, NaCl, citric acid, or quinine; 1/day) for 30 min, and consumption was measured by weight. A subsequent one-way ANOVA revealed strong inter-stimulus differences in consumption (F(3,28) = 32.97, *p* < 0.01)— the order of taste preferences was sucrose>NaCl>citric acid>quinine.

We then, for each window of neural firing (window size: 250 ms; step size: 25 ms), performed a Spearman product-moment correlation between the ranked firing rates to each taste and the palatability rankings (Spearman was used, rather than the parametric Pearson, because palatability rankings are ordinal variables). Note that, for this analysis, it matters not to what degree an individual taste was or was not preferred to water; the calculated correlation, like the 2-way ANOVA used to measure taste-specificity of firing, cleanly measures the palatability-relatedness of firing by relating the activity driven by each taste to that driven by the others. But because this procedure is another that results in a relatively large number of actual tests, we limited the chances of Type I errors by requiring that the test achieve significance for 3 consecutive bins before responses could be deemed palatability-related.

### Modeling of, and firing-rate change-point identification in, ensemble firing data

GC taste processing is dynamic—rat cortical responses have been shown to undergo a pair of sudden and coherent firing-rate transitions in the 1.5 sec immediately following taste delivery; the state prior to the first transition is the same for all tastes, reflecting taste ‘detection;’ the states attained following the first and second transitions are both taste-specific, with the last state further reflecting taste palatability (and both predicting and driving taste behavior in single trials, see Sadacca et al., 2016; Li et al., 2016). In order to detect whether such transitions between states are also facets of mouse cortical taste processing, we performed a simple change-point analysis on ensemble spike data. This analysis is summarized here; for complete details, see Mukherjee et al. (2017).

Trials of ensemble spiking data were categorized in terms of which neuron spiked in each 1-ms bin; an index of 0 corresponded to no spikes from any neuron. If more than one neuron spiked in a time bin—a highly uncommon occurrence, given the relatively low firing rates of GC neurons—we randomly selected one spiking neuron to index that bin (Jones et al., 2007; Sadacca et al., 2016). We then fit a change-point model to the first 1.7s of post taste delivery ensemble activity, describing ensemble firing rate transitions between states as categorical distributions with emissions. We initially considered the possibility that our mouse GC data might be best fit by models containing fewer or more than 2 change points. To compare the quality of the fits of the different models to the ensemble spiking data, we computed the Akaike information criterion (AIC) using the formula AIC = 2*k - 2*LL, wherein k represents number of parameters and LL is log likelihood of a particular model, and then examined the properties of the identified change points.

As the results of this analysis confirmed the appropriateness of the 3-state (i.e., 2-change point) model (see Results), we constructed a more extensive model constrained on the basis of the rat cortical findings described above: specifically, we constrained the change from state 1 to state 2 (change-point 1, or CP1) to happen within the interval [50 – 600 ms], and constrained the second change-point (CP2) to occur within the interval [CP1 + 0.2s to 1500ms]. This is equivalent to placing uniform priors over the intervals that define CP1 and CP2, corresponding to the timing of sudden, coherent firing rate transitions in GC ensembles (Jones et al., 2007, Sadacca et al., 2016). With these assumptions, the Expectation-Maximization (EM) algorithm, the most widely used approach to perform inference in such models with latent variables (CP1 and CP2), was employed to fit the change-point model (Bishop, 2016).

Note that to facilitate a central test of this model, the above-described constraints on the timing of state changes are broad and deliberately not centered on the average timing of the transitions as previously observed in rats: specifically, on the basis of our previous work, we predicted CP1 to occur on average at ~200 ms post-delivery, and predicted CP2 to occur on average at ~900 ms post-delivery. If ensemble spiking fails to go through genuinely sudden, coherent firing transitions, any change points identified by the analysis would instead appear (on average) in the middle of the available interval. This result, or any change-point timings, would falsify the prediction that mouse GC ensemble taste responses resemble rat GC ensemble taste data.

### Histology

At the end of each experiment, mice were deeply anesthetized and perfused transcardially with 0.9 % saline for 1 minute followed by 4% paraformaldehyde. Brains were extracted and post fixed in 4% paraformaldehyde for 24-48 hours, after which coronal brain slices (60 μm) containing the GC were sectioned on a vibratome. Sections were rinsed 5-6 times over 90 mins (PBS/.3% Triton X-100), mounted on charged glass slides, and cover slipped with mounting medium including DAPI (Vectashield). The placement of electrodes was determined by localizing Dil (a lipophilic dye coated on electrodes during implantation). To visualize electrode bundle locations, bilateral GC sections were viewed by confocal fluorescence microscopy with a Leica Sp5 Spectral confocal microscope/Resonant Scanner with 405 (for DAPI), 546 (Dil dye) and 488 (for GFP) lasers equipped with x/y/z movement stage.

## Results

A total of 185 units were isolated across 10 sessions in 6 mice (10-37 neurons/ensemble)—4 mice received 2 sessions each (in which case sessions were recorded from separate DV locations in the GC; see material and methods). Our recordings spanned much of the GC region (AP Bregma +1.6 to 0.0; DV 2.25 – 3.25 mm from the surface of the brain; see Figure 1) receiving input from taste thalamus (Chen et al., 2011; Fletcher et al., 2017) or from the amygdala (Matyas et al., 2014). Prior work has shown these regions to be important for taste behavior and learning in mice (Neseliler et al., 2011).

### Basic characterization of taste evoked responses in the gustatory cortex of mice

Figure 2 presents raster plots and associated PSTHs of the taste responses of three simultaneously recorded GC neurons. The first two are regularly spiking (putative) pyramidal neurons and the third is a fast-spiking (putative) inhibitory interneuron. Note that we did not rigorously identify every recorded neuron as a (putative) pyramidal cell or interneuron using baseline firing rates and action potential widths here; we have done so previously, and this prior work revealed no evidence of major cell-type-specificity with regard to any of the analyses performed below (Fontanini and Katz, 2006; Jones et al., 2007; Katz et al., 2001).

**Figure 2.**
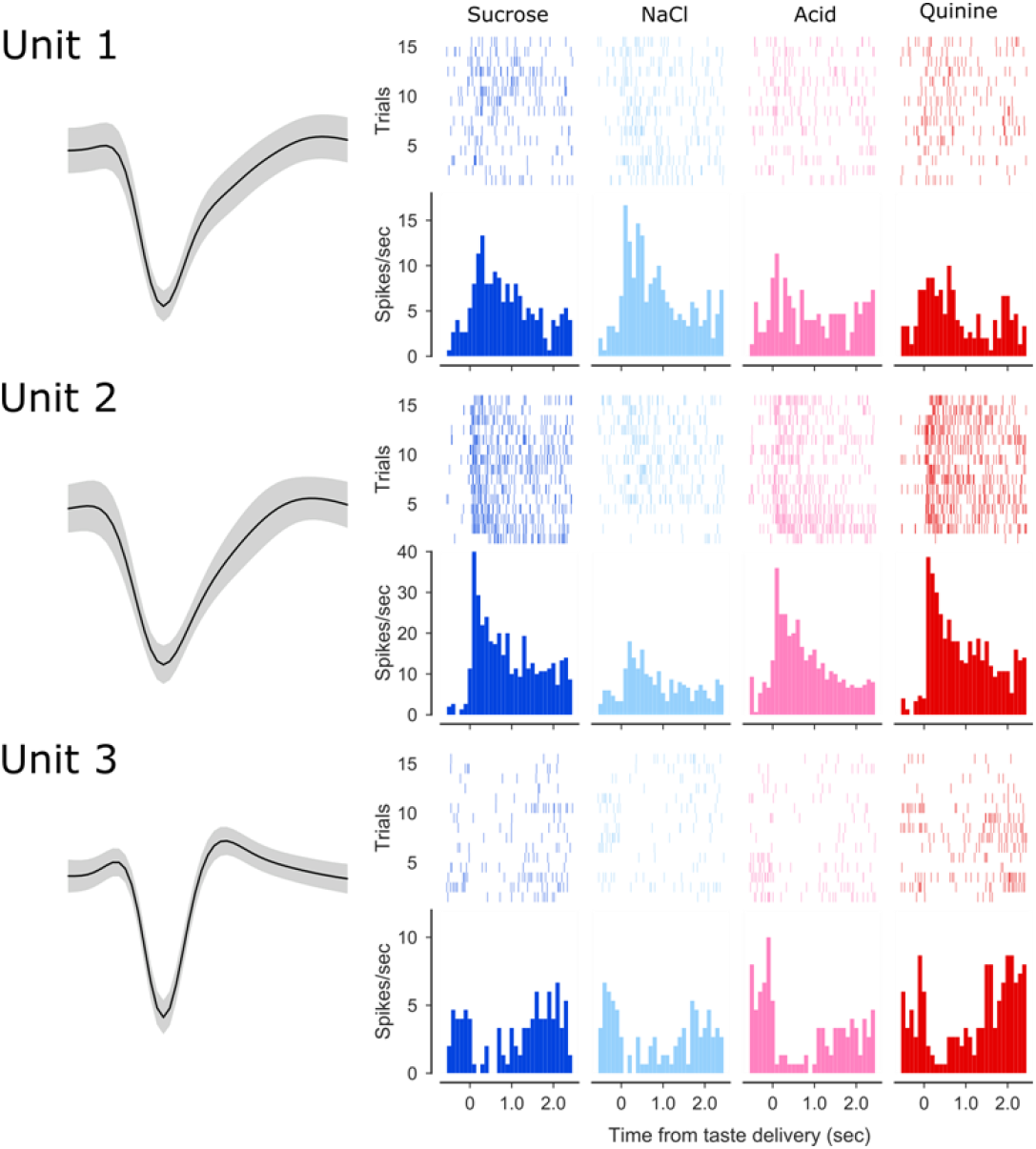
Simultaneously recorded gustatory cortex neurons. The left column shows the average action potential waveforms (surrounded by confidence intervals) for three neurons recorded simultaneously from the same electrode bundle. The remaining four columns show the responses of these neurons to sweet (sucrose), salty (sodium chloride), sour (citric acid) and bitter (quinine) tastes. The top half of each response display is spike time raster plots (each row is a single trial, and each dot is an action potential), below which are associated PSTHs (firing rate in spike/second in the y-axes; time post-stimulus in the x-axes). Each neuron’s responses were taste-specific (see Methods), and each individual neuron responded distinctly.

Each of these neurons’ responses were significantly (see Methods) taste-specific: Unit 1 responded to all 4 tastes, but responded more strongly to NaCl and sucrose than to the remaining tastes; Unit 2, on the other hand, responded strongly to sucrose, citric acid, and quinine, but far less to NaCl; Unit 3, meanwhile, was inhibited to differing degrees and for differing lengths of interval by all 4 tastes.

Even the most basic examination of the full sample of responses revealed that a large percentage (just under 2/3) of mouse GC neurons respond to taste stimulation (significant paired t-tests comparing pre-to post-stimulus firing across all trials summed across tastes, Figure 3A), with excitatory responses (enhanced firing) outnumbered inhibitory (reduced firing) by almost 3 to 1. Using repeated-measures ANOVAs to directly compare each neuron’s average and time-varying response to different tastes, we were able to identify an almost identical percentage of neurons as being “taste neurons” (i.e., responding distinctly to at least one taste, see Methods and Figure 3C). While these particular t-tests and ANOVAs will not necessarily identify precisely the same neurons, together they suggest that the vast majority of “taste responses” in mouse GC are truly taste-related, as opposed to reflecting somatosensory or general cognitive factors.

**Figure 3.**
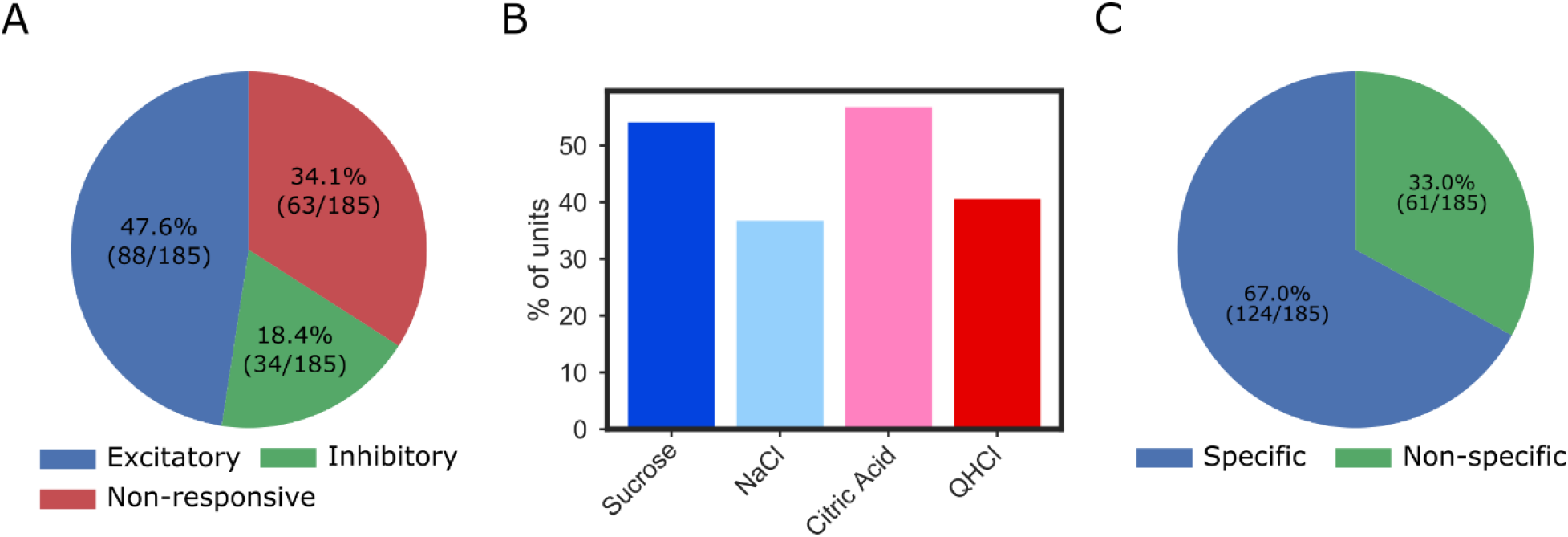
Mouse GC neurons respond to tastes. **A**. A pie chart showing the percentage and distribution of GC neurons that produced excitatory or inhibitory responses to taste stimulation. **B**. Similar percentages of mouse GC neurons respond to sweet, salty, sour, and bitter tastes; at least 35% of the sample responded to each taste. **C**. 2-way within-neuron ANOVAs comparing responses to the different tastes as a function of time (taste x time, *p*<0.01) revealed that the majority of mouse GC neuron taste responses were also taste-specific (i.e., different for different tastes).

Despite being taste-specific, these responses were broad—significant firing rate modulations in individual neurons were seldom restricted to only one, or even two, of the four basic tastes delivered. Had they been narrow, then the rate of significant responses to each individual taste (determined using simple t-tests, as above, applied to each individual PSTH) would be expected to be around ¼ of this 66% (that is, we’d expect ~16% of the neurons to have sucrose responses, another ~16% to have NaCl responses, etc). A far higher percentage of neurons responded to each taste, however (Figure 3B), a result consistent with the suggestion that most taste neurons in mouse GC must respond to > 1 taste on average.

Because this sort of analysis carries with it a host of (commonly ignored) caveats (see Methods), we performed multiple convergent tests of this conclusion. First, we directly analyzed the number of tastes that evoked significant responses in each individual neuron (by pre-vs post-stimulus t-tests). According to this analysis, 28.6% of our mouse GC neurons responded significantly to one taste, 25.6% responded to two tastes, 16.8% to three tastes, and 14.1% to all four tastes (Figure 4A). While it is inappropriate to reach conclusions concerning taste specificity from this method (which does not directly compare responses to different tastes, see Methods), these results confirm that the majority of GC neurons that respond to taste stimulation respond to > 1 taste.

**Figure 4.**
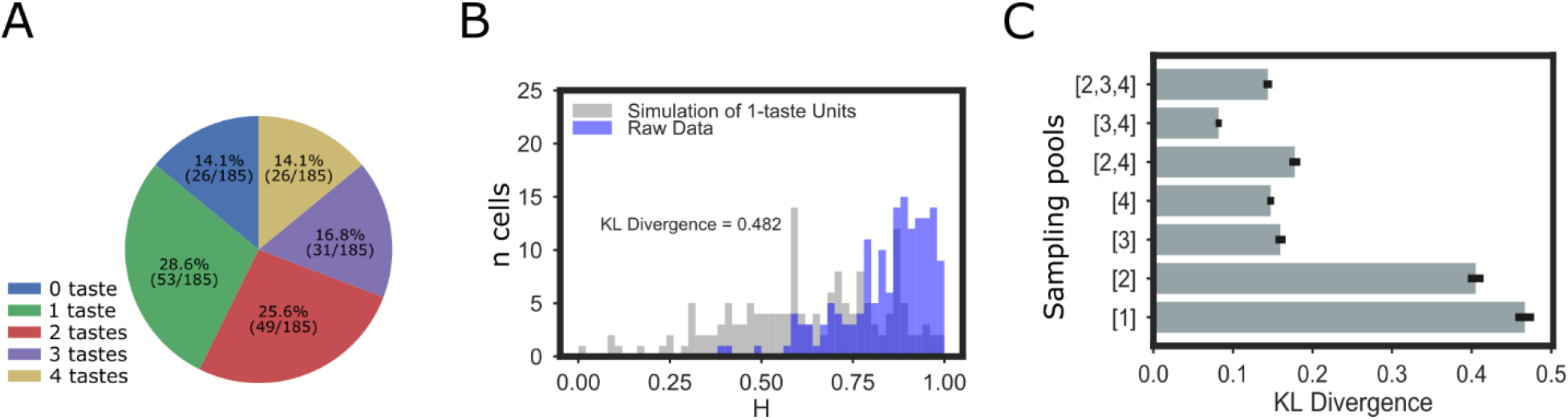
GC neurons code tastes broadly. **A.** Independent paired t-tests comparing pre- and post-taste firing rates reveal that the majority of mouse GC neurons respond more than one taste (even when totally non-responsive neurons are included in the comparison). **B.** The distribution of the breadth of taste coding (i.e., entropy [H], x-axis) in GC neurons (purple bars, y-axis = 159 of neurons) is highly skewed toward broad tuning (a low H indicates narrowly responsive neuron and a high H indicates a neuron responsive to several tastes). Overlain (grey bars) is the distribution of response entropies for an equivalently-sized set of simulated units which respond to only one taste. The difference of the two distributions was calculated using the Kullback-Leibler (KL) divergence, and was highly significant according to a *X^2^* test conducted on the data. **C**. The KL divergences between the collected data and a range of simulated datasets in which taste responses were simulated from units sampled from the population of neurons responding to particular numbers of tastes (sampling pools).

We went on to calculate response entropy (H) for each neuron, using standard techniques (Smith and Travers., 1979; Samuelsen et al., 2013); low H values indicate narrowly responsive neurons, and high H values indicate broadly tuned neurons. The distribution for our neural sample proved to be highly positively skewed, with the vast majority of GC neurons’ H values suggesting broad responsiveness; the distribution was similar when only neurons identified as responding in a taste-specific manner were included (Figure 4B).

To enhance the interpretability of this last result (see Methods), we plotted (Figure 4B) the distribution of entropies against the H values of a simulated set of neurons modeled on those real neurons identified as responsive to a single taste. This distribution, indicated by Chi-squared analysis, is significantly different from the distribution of the full set of real data (*X^2^* = 82.75, *p* < 0.01), which, in fact, most resembles (according to the Kullback-Leibler divergence) simulated data sets modeled on neurons that respond to 3 or 4 tastes (Figure 4C).

This analysis also reveals that even neurons that in Figure 4A are listed as responsive to only 1 taste are not 100% unresponsive to other tastes. Was this not the case—that is, was neural firing totally unaffected by administration of 3 out of the 4 tastes, the calculated entropy would equal 0.0; instead, many “1-taste” neurons actually produced sub-significant firing-rate changes to multiple other tastes, a fact that leads even these “1-taste” neurons to show entropy values greater than 0.5 (see detailed discussion in Smith and Travers, 1979). Thus, Figures 4B and 4C are consistent with the suggestion that mouse GC neurons tend toward broad tuning.

### Dynamics of taste-evoked responses in mice GC

In revealing GC taste neurons to be broadly tuned “coarse coders,” the above data are consistent both with previous reports of taste responses in rat GC (Yamamoto et al., 1985; Bahar et al., 2004; Fontanini and Katz., 2006; Jones et al., 2007; Piette et al., 2012; Samuelsen et al., 2013), and with calcium imaging data from mouse GC (Fletcher et al., 2017; although see Chen et al., 2011). But further examination reveals that these above analyses, and the conclusions drawn from them, inadequately describe taste processing in mouse GC, in that they ignore meaningful response dynamics that unfold across the first 1.5s of post-stimulus time (the period leading up to the making of consumption decisions and production of consumption-related behaviors, see Grill & Norgren, 1978; Travers and Norgren., 1986; Li et al., 2016)—response dynamics that can only be reliably observed using multiple trials of electrophysiological data.

Even simple visual scrutiny of mouse GC taste PSTHs suggests the presence of such interpretable firing-rate dynamics, with response magnitudes changing through time in taste-specific manners. An example is shown in Figure 5A—a mouse GC neuron that responded strongly to NaCl during the 1^st^ post-delivery second only, and strongly to quinine during the 2^nd^ post-delivery second only. When this pattern of firing is averaged across time, both of these responses “wash out,” and the neuron incorrectly registers as being responsive to citric acid only; this result confirms the need to move beyond analyses that average across time, if the purpose is to comprehensively characterize taste-specificity of firing in periods leading up to behavioral responses (see Discussion).

**Figure 5.**
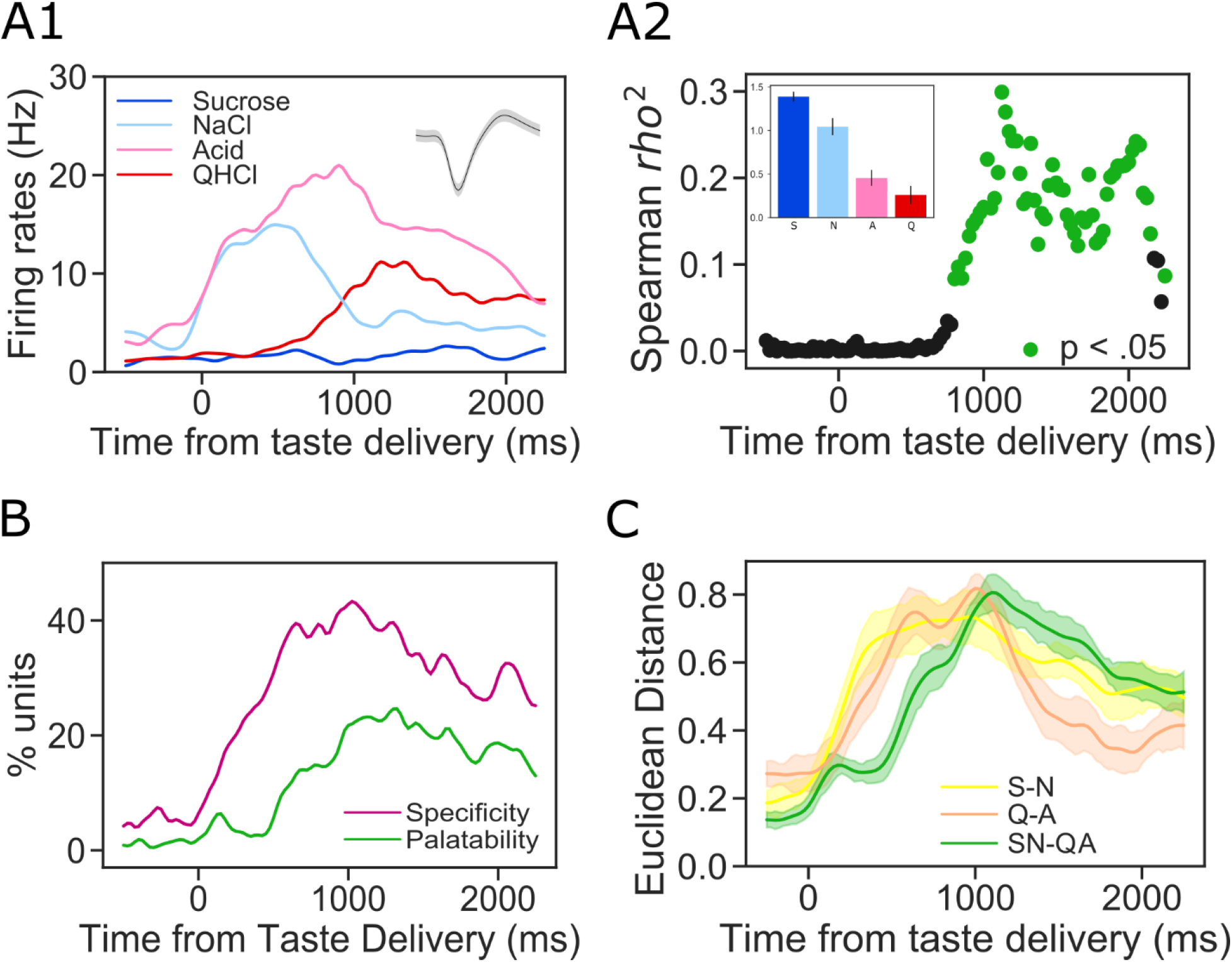
Mouse GC responses evolve through successive epochs of taste quality and taste palatability coding. **A1**: Overlain PSTHs (y-axis, firing rate; x-axis, time post-stimulus) from a representative GC neuron that responded significantly to all tastes but sucrose [S]. **A2**: Moving window analysis of the correlation (R-squared, y-axis) between firing rates and taste palatability (determined by taste intake, see inset) for the neuron shown in Panel A1. Green dots denote significantly non-zero correlations. **B**. Percentages of the entire sample of GC responses that show significant taste specificity (magenta) and palatability-relatedness (green). Taste specificity starts to rise soon after taste delivery, and peaks at 500ms; in contrast, palatability-related firing appears only following a delay (~500ms), and peaks at ~1 second post-stimulation. **C**. Response differences (Euclidean separation in multi-dimensional scaling solutions, see methods for details) between same-palatability tastes qualities (S-N and Q-A) emerge soon after stimulus delivery; between-palatability differences, on the other hand (SN-QA) emerge later, confirming that identity-related activity is followed by palatability-related firing.

Previous work using rats has suggested that an intrinsic part of GC taste response dynamics is the relatively late transition from taste-specific firing into palatability-related firing. To rigorously evaluate the hypothesis that mouse GC taste responses start taste-specific and then become palatability-related, we first ascertained behavioral preferences for our battery of four tastes (which differed in both chemical identity and palatability, sucrose and sodium-chloride representing palatable tastes and citric acid and quinine representing non-palatable tastes) in the same strain of mice being used in the electrophysiology experiments, using a single-bottle consumption test (see Methods). We then correlated this vector of relative palatabilities (note that we are concerned here, as we were when evaluating taste-specificity of firing above, with differences between tastes, and not with absolute levels of “palatability”) with firing rates to the same battery of tastes. For this purpose, firing was calculated for small bins across the 2-sec response epoch (using a moving-window analysis as explained in the Methods and used previously in Fontanini et al., 2009; Piette et al., 2012; Sadacca et al., 2012; Li et al., 2016; Sadacca et al., 2016).

The result of this analysis brought to bear on the neuron shown in Figure 5A1 is shown in Figure 5A2 (with the empirical taste preference data presented as an inset): this single neuron fired in a taste-specific manner almost from the moment of taste delivery, but firing did not become significantly palatability-related (green dots) until several hundred msec later, and palatability-relatedness only reached an asymptote a full sec after taste delivery. In this case the ordering was “aversive high,” which appeared with approximately the same frequency in our neural sample as “aversive low.” As described above, this pattern reflects that fact that firing rates were modulated at least twice in succession following taste delivery—the first firing-rate modulation resulted in taste-specificity, and the second in palatability-relatedness.

Figure 5B, which presents the results of this analysis performed upon the entire data set, confirms the representativeness of the example: the incidence of taste specificity rises almost immediately following taste stimulus delivery, and approaches peak levels as early as 500 ms into the response; palatability relatedness, on the other hand, rises more slowly, beginning to trend upward ~500 ms after stimulus delivery and approaching a peak ~1000 ms into the response.

We performed one further test of this result, analyzing the distinctiveness of different subsets of taste responses for each neuron at each point in response time; “distinctiveness” was quantified in terms of normalized Euclidean distances separating two taste responses in a multi-dimensional scaling solution. Given the above results, we predicted that the distinctiveness of individual pairs of tastes would rise quickly, but that the distinctiveness of palatability differences—that is, comparisons of the averaged responses to the palatable tastes (sucrose and NaCl) to the averaged responses to the aversive tastes (quinine and citric acid)—would emerge only later. Figure 5C reveals that this is precisely what we observed: the distinctiveness of codes for sucrose and NaCl (yellow line ± SE) rose quickly in post-delivery time, as did the distinctiveness of codes for quinine and citric acid (beige line ± SE); the rise in the distinctiveness of palatabilities (green line ± SE), meanwhile, was delayed. Note as well that following the rise of palatability distinctiveness, the distinctiveness of individual “same-palatability” taste pairs fell precipitously, confirming that general taste-specificity becomes refined across time into palatability-relatedness.

Thus, we conclude that mouse GC codes taste properties sequentially across 1.5 seconds of post-stimulus time, with taste-specificity appearing prior to palatability-relatedness. Ancillary analysis confirms that many individual neurons respond distinctly in the two epochs, coding both taste identity early and taste palatability late. In fact, 50% of the neurons that produce significantly taste-specific firing also produce significantly palatability-related firing (as in the example neuron shown in Figure 5A). This result is novel, and in fact can currently only be recognized using electrophysiology—the subtle difference in responses that we report, and shifts in those differences across a few hundred milliseconds, are almost impossible to discern with calcium imaging data.

### Ensemble properties of mouse cortical taste dynamics

Having established the existence of reliable, interpretable single-neuron dynamics in mouse GC, we moved on to asking whether these single-neuron dynamics are in fact reflections of coherent ensemble activity (as they are in rats, see Sadacca et al., 2016). To answer this question, we applied multi-signal change-point (CP) modeling procedures (Mukherjee et al., 2017) to datasets comprised of simultaneously-recorded neural ensembles, to test whether transitions between the above-observed firing-rate epochs are: 1) coherent between neurons; 2) sudden and jump-like; and 3) trial-specific in latency.

We first compared our ability to model the data using 2 CPs (the predicted model given the above results and previous work on rats) to that achieved using models with fewer or more CPs, with the logic being that if our hypotheses are correct, a 2-transition model should fit the data particularly well. Initial assessment of the quality of model fit making use of the Akaike information criterion (AIC) confirmed that mouse GC ensemble taste data are typically well characterized as going through a set of three states separated by two CPs (2 sudden transitions of firing that were coherent across simultaneously-recorded neural ensembles). While in a subset of instances a 3-CP model fit equally well, scrutiny of these successful 3-CP fits revealed that, in over half of the trials, multiple CPs were clustered with minimal time separations—they were essentially 2-CP solutions with one of the CPs identified twice. Similar results were observed in a subset of datasets in which the fit of a 4-CP model was examined. Clearly, mouse GC taste responses are best described as going through a sequence of 3 firing-rate “states” separated by sudden, coherent ensemble firing-rate transitions.

Our analysis of the data as characterized by the 2-CP model demonstrates further features of the taste code in mouse GC. Figure 6A shows an example of 3 QHCl trials showing within-trial firing rate transitions. Clearly, the transitions happen at different times on different trials. To test whether the transitions relate to the earlier-described epochs, we once again correlated palatability with firing, but did so separately for the data in each successive state (putting all “State 2” firing into one analysis, e.g.). T he results of these analyses are displayed in Figure 6B; a one-way ANOVA revealed that the palatability correlation varied with state, *F(2, 368)* = 42.80, *p* < 0.01, and post-hoc comparisons (*Tukey HSD* test) further revealed that this significance is largely driven by the fact that the correlation with palatability is significantly higher following the second transition than before (*p* < .05). The fact that palatability correlations are modestly higher following the first transition than before reflects the simple fact that the very earliest 100-200ms of GC responses (i.e., the brief period preceding even the 1^st^ transition) are, as they are in rats, totally non-specific; almost any taste specificity will enhance correlations with palatability beyond those chance correlations observed with non-specific firing.

**Figure 6.**
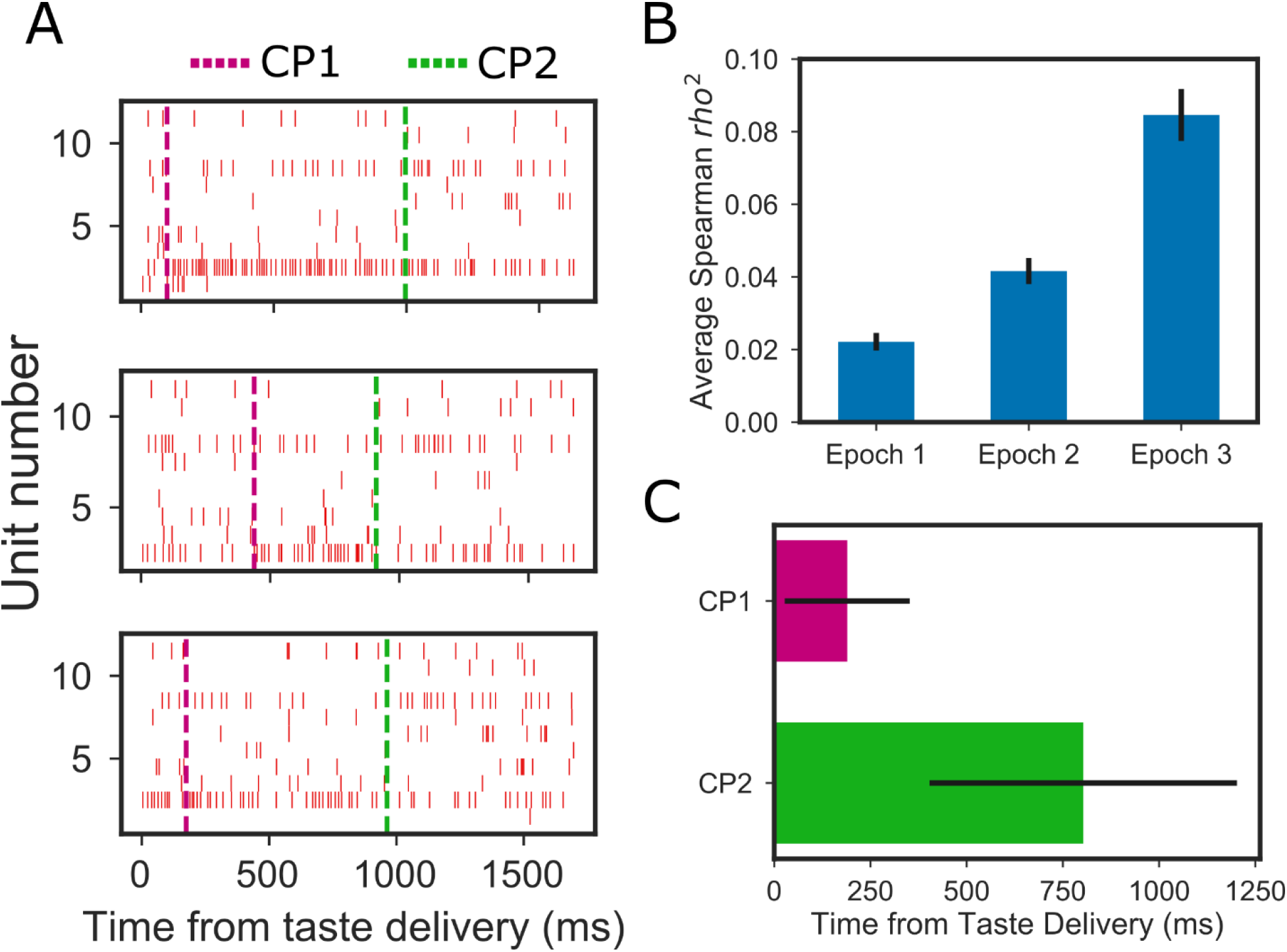
The epochs of mouse cortical taste responses reflect coherent state sequences in GC ensembles. **A**. Representative single trials of the response of one GC ensemble (one neuron/row; hash marks = individual action potentials) to the taste of QHCl, revealing sudden, simultaneous changes in firing rates in several neurons. Magenta and green dashed lines indicate the first switch point (CP1: from taste detection to identity) and second switch point (CP2: from taste identity to palatability), respectively. **B**. Coefficient of determination (r-squares) between mean firing rates and palatability for each of the three states, demonstrate, consistent with our hypotheses and work on rat GC, that palatability coding is achieved with state 3, and that taste-specificity (which necessarily connotes a basic level of non-zero palatability coding) is achieved with state 2. **C**. Average state-to-state transition times across all trials revealed through switch point analysis.

The average transition times for CP1 and CP2 occurred at ~190 ms and ~800 ms, respectively (Figure 6C). Note that these average transition times differed distinctly from the time-point that represented the middle of the available intervals (i.e., 325 for CP1 and 925ms for CP2; see Methods). This finding proves that the CPs are not simply artifacts of the procedure itself; rather, these latencies are good matches for our prior rat data, and for our single-neuron results (Fig 5D), and confirm our expectation that mouse GC ensembles undergo fairly sudden and coherent shifts in firing while responding to tastes.

These results suggest that even dynamic analysis of single-neuron taste responses in mouse GC, as described above, fail to completely characterize taste processing. Rather than being independent coding elements, single-neuron responses are likely reflections of population coding—an unsurprising fact given the mesoscopic (intracortical connectivity) and macro-scopic (between-region feedback) circuitry involved in taste, but one that has not been recognized previously.

### Spatial analysis of mouse cortical taste response properties

GC is a distributed, heterogeneous region that can be subdivided according to cytoarchitectural and connectivity criteria in the dorsal-ventral and anterior-posterior axes (Allen et al., 1991; Maffei et al., 2012). The imaging literature has failed to provide clarity on the significance of these anatomical subdivisions—one set of authors has suggested that mouse cortical taste coding, unlike that of rat (Accolla et al., 2007), is cleanly gustotopic (Chen et al., 2011; Wang et al., 2018), while other research groups suggest otherwise (Fletcher et al., 2017; Lavi et al., 2018).

To explore the potential spatial dependencies of GC taste coding using electrophysiology, we compared taste-evoked firing along the two primary axes of GC. In the dorsal-ventral dimension we were able to look within-subject, as our electrode bundles could be driven ventrally between tasting sessions. Figure 7 shows a pair of example neurons recorded from dorsal GC, and a second pair recorded from ventral GC in the same mouse. It can be seen that each of these responses are quite broad, suggesting basic similarities between dorsal and ventral GC.

**Figure 7.**
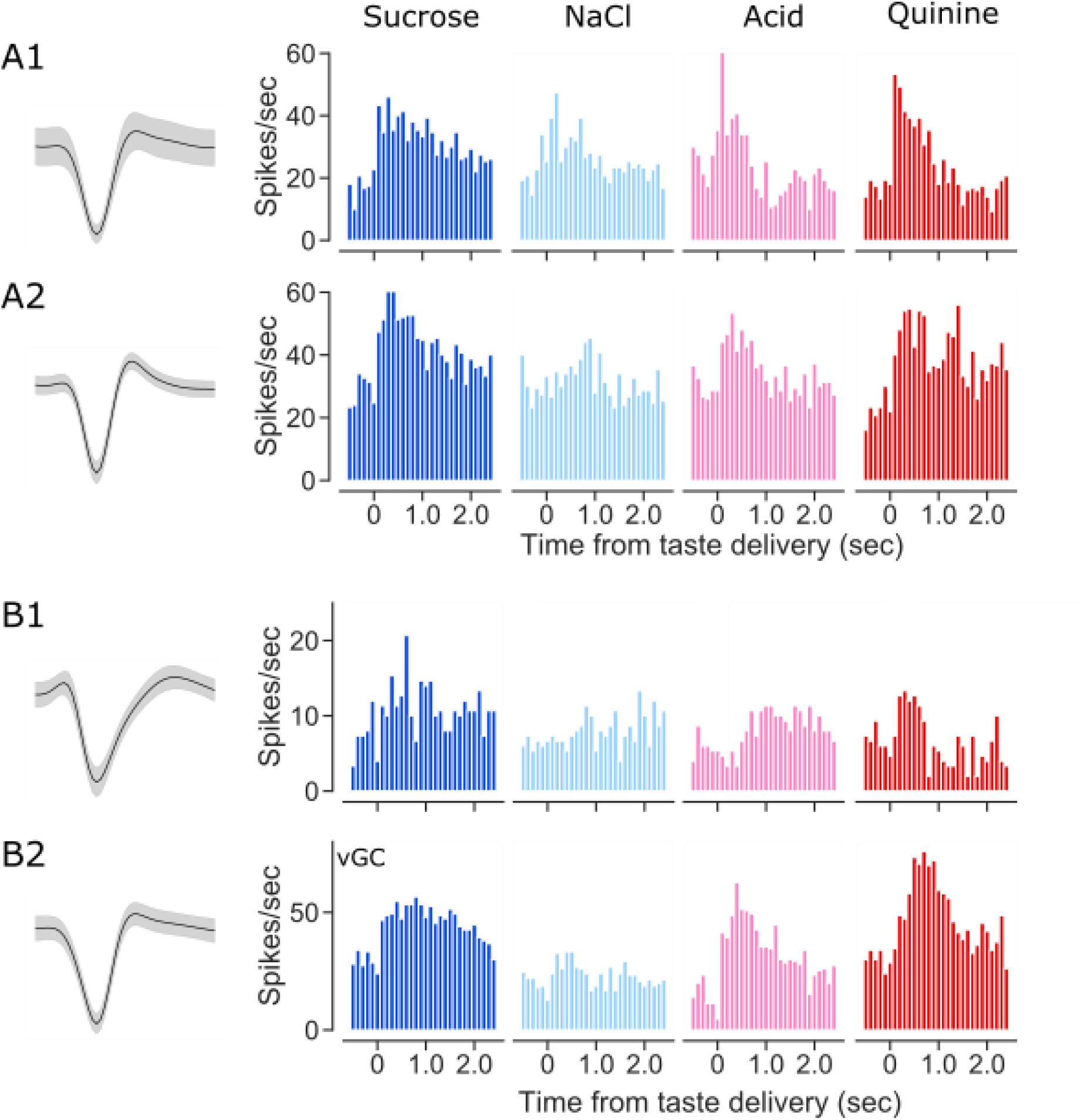
Broadly-responsive ensembles of GC neurons can be recorded from dorsal (A1, A2) and ventral (B1, B2) GC of the same mouse. **A1-A2**. Representative neurons (wave shapes and PTSHs, conventions as for Figure 2) recorded in dorsal GC. **B1-B2**. Representative neurons recorded from the same mouse in a 2^nd^ session.

Closer analysis confirmed that both dorsal and ventral GC contained neurons that produced broad, dynamic taste responses. Analysis of the entire sample divided into dorsal and ventral recordings revealed no striking differences in breadth of responsiveness, measured either in terms of number of tastes responded to (Figure 8A, *X^2^* = 1.15, *p* = 0.885) or response entropy (Figure 8B; *X^2^* < 1). The magnitudes of taste specificity revealed by ANOVA was similar across dorsal and ventral neurons (Fig. 8C; *F* < 1), and the pattern of how many neurons responded to which taste did not differ significantly between subregion (Figure 8D); in both dorsal and ventral GC, single neurons were more likely to respond to both palatable and aversive neurons than either alone (data not shown). Finally, dorsal and ventral neural samples showed similar time-courses of response evolution from quality to palatability, measured in terms of the number of neurons showing such responses at each time point (Figure 8E1-2).

**Figure 8.**
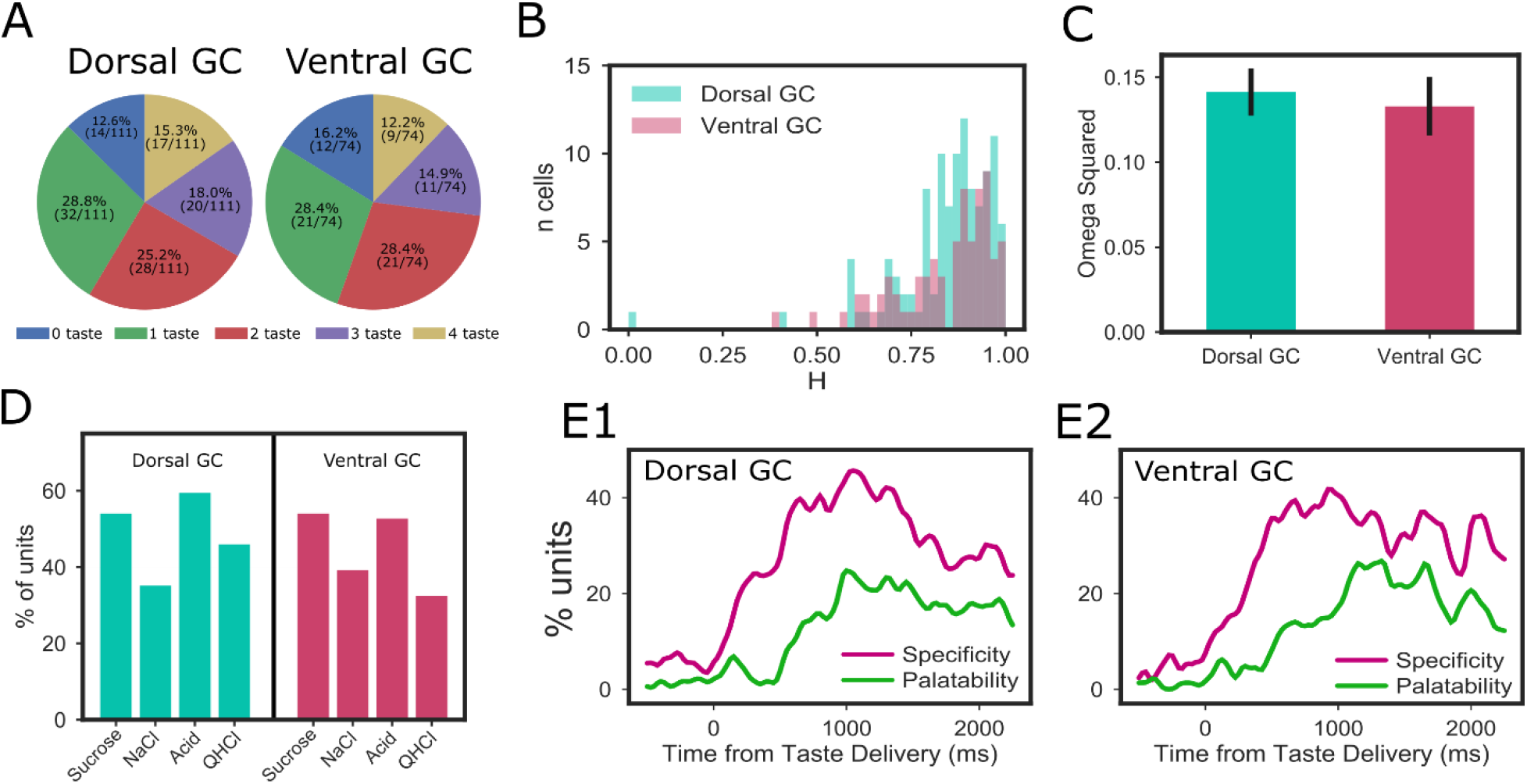
Dorsal and ventral GC taste responses in mice have similar properties. **A**. Percentages of dorsal and ventral neurons with significant responses to zero, one, two, three, and four tastes. X^2^ tests revealed no significant differences between the distributions. **B**. Response entropies (H) for dorsal and ventral GC neurons were not significantly different, according to X^2^ tests comparing the distributions. **C**. The magnitudes of taste specificity (expressed as omega squared from 2-way ANOVAs, see Methods for details) for dorsal and ventral GC neurons. T-test revealed no difference. **D**. The distribution of relative responsiveness to the four tastes is virtually identical in dorsal and ventral GC—a X^2^ test did not suggest any differences in which subregion responded to which tastes. **E1-2**. In both dorsal and ventral GC, taste specificity starts to rise immediately after taste delivery and peaks at 500ms, and in both, the emergence and peaking of palatability-related firing was delayed (see Figure 5).

The results of our analysis of the anterior-posterior dimension were analogous. We compared taste responses between bregma +1.6 and bregma 0.0, treating everything forward of 0.8 as “anterior.” This analysis revealed no significant differences between anterior/posterior breadth of responsiveness, measured either in terms of number of tastes responded to (Figure 9A, *X^2^* = 5.51, *p* = 0.239) or response entropy (Figure 9B; *X^2^* < 1); furthermore, neurons from anterior and posterior GC subdivisions had similar magnitudes of taste specificity (Figure 9C; *F* < 1), similar patterns of how many neurons responded to each taste (Figure 9D), and similar time-courses of response evolution from quality to palatability (Figure 9E1-2).

**Figure 9.**
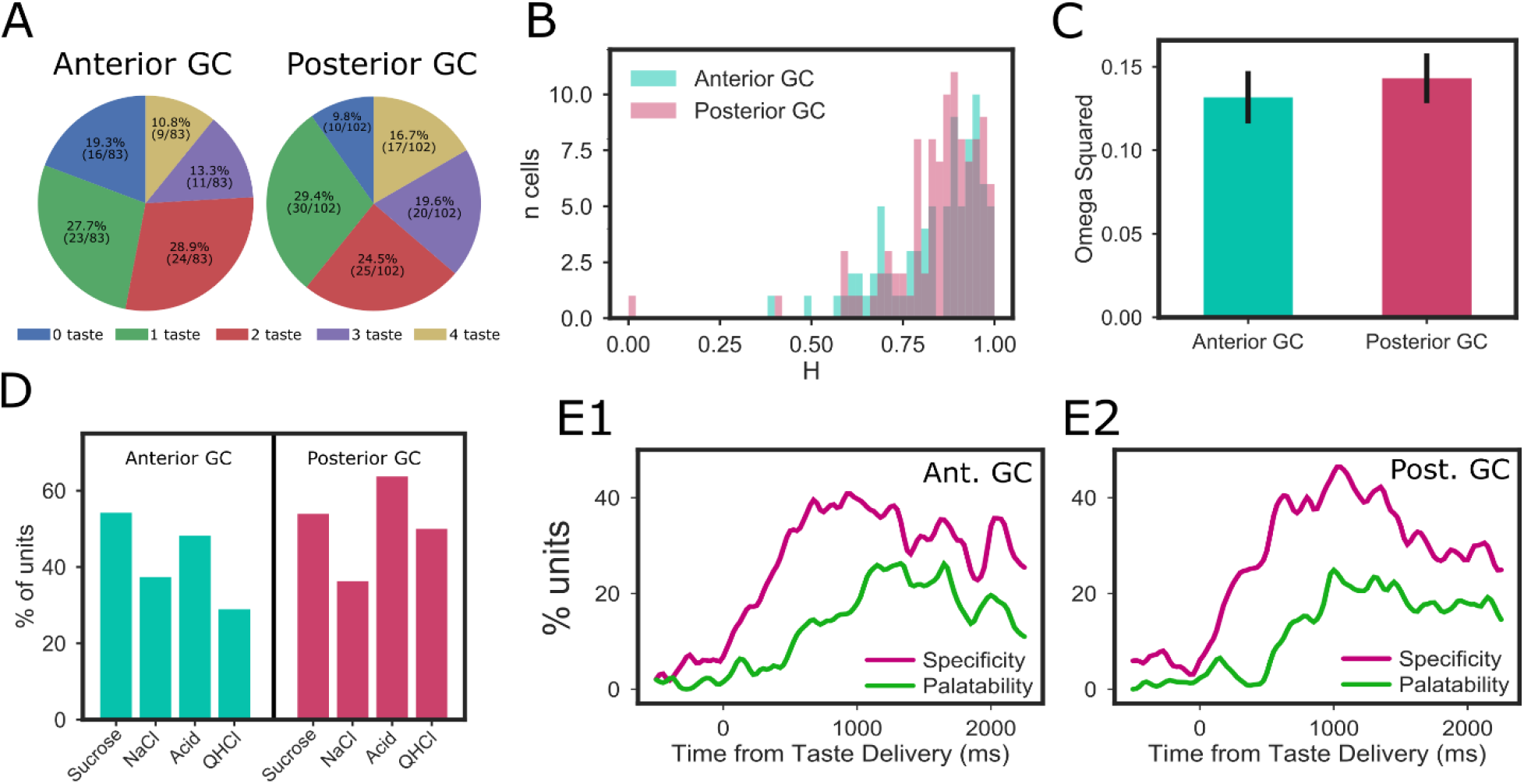
Anterior and posterior GC taste responses in mice have similar properties. **A**. Percentages of anterior and posterior neurons with significant responses to zero, one, two, three, and four tastes. X^2^ tests revealed no significant differences between the distributions. **B**. Response entropies (H) for anterior and posterior GC neurons were not significantly different, according to X^2^ tests comparing the distributions. **C**. The magnitudes of taste specificity (expressed as omega squared from 2-way ANOVAs, see Methods for details) for anterior and posterior GC neurons. T-test revealed no difference. **D**. The distribution of relative responsiveness to the four tastes in posterior GC was similar to that in anterior GC; while responsiveness to the two more aversive tastes was somewhat higher in posterior GC, a X^2^ test did not reveal any significant differences in which subregion responded to which tastes. **E1-2**. In both anterior and posterior GC, taste specificity starts to rise immediately after taste delivery and peaks at 500ms, and in both, the emergence and peaking of palatability-related firing was delayed (see Figure 5).

Together these results suggest that the details of taste coding are relatively insensitive to anatomical location within mouse GC. Neurons across the anterior-posterior and dorsal-ventral axes code taste modalities in a broad and temporally complex manner. Of course, the lack of significant difference between anatomical sub-regions does not equal rigorously identified “sameness,” and the possibility remains that subtle differences between anatomical subregions (or between species, for that matter) remain to be identified. Nonetheless, these analyses lead us to conclude that whatever differentiates different parts of gustatory insular cortex (see Discussion), it is neither the breadth nor dynamics of single-neuron taste responsiveness.

## Discussion

Mice have become the most common non-human mammalian species studied by neuroscientists—a fact that can no doubt be attributed to the accessibility of mouse genetics, which allows researchers to, with (relative) ease, study the underlying molecular mechanisms of cellular, network, and behavioral phenomena (Stevens, 1996; Kandel et al., 2014). As the sense with (arguably) the tightest link to behavior and learning (Carleton et al., 2010; Maffei et al., 2012; Katz & Sadacca, 2010), taste is a particularly good system with which to study these topics in a unified manner. Furthermore, this work can take advantage of the extensive progress that has been made toward understanding key principles of taste coding in awake rats (Accolla et al., 2007; Bahar et al., 2004; Fontanini and Katz., 2006; Jones et al., 2007; Katz et al., 2001; Li et al., 2016; Moran & Katz., 2014; Sadacca et al., 2012; Samuelsen et al., 2012). It is somewhat surprising, therefore, that there has been almost no electrophysiological work done on mouse cortical taste coding, and that the imaging studies performed thus far have failed to provide consensus on basic features of gustatory sensory coding (Chen et al., 2011; Fletcher et al., 2017; Lavi et al., 2018).

In fact, while electrophysiological responses to stimuli including tastes have been examined in anesthetized mice (Wilson and Lemon, 2013), the current study represents the first thorough investigation of single-neuron spiking responses to a broad battery of taste stimuli by primary gustatory cortical (GC) neurons in awake mice (but see Kusamoto-Yoshida et al, 2015). By delivering a battery of four basic tastes (sweet, salty, sour and bitter) directly to the tongue of freely moving mice while making recordings from a multi-electrode bundle designed to record the activity of small ensembles of single neurons, we are able to confirm the conclusion reached by Fletcher et al (2017) and Livneh et al (2017)—namely, that mouse GC codes taste in a largely non-sparse manner: approximately 2/3 of recorded GC responses conveyed information about taste identity; by and large, these responses were broadly tuned—over half of the recorded neurons responded to > 1 taste, and over half of taste neurons produced firing patterns that were distinctive for 3 of 4 tastes.

But we were able to go further, to reveal aspects of mouse GC taste responses that are largely hidden in calcium imaging data. As detailed in the Methods section, the fine-grained temporal resolution and high signal-to-noise ratio offered by electrophysiology allowed us to show that the time courses of taste responses contain information about the taste (see Figure 5A for an example)—that between-taste differences in time course are not random. Specifically, population ensemble analysis (Figure 6) shows that the mouse GC taste responses evolve in a characterizable way across 1-1.5 seconds. We identify separate “epochs” of GC responses related to the physiochemical and psychological properties of the tastes in our stimulus battery. A large portion (50%) of the recorded neurons coded stimulus identity and stimulus palatability sequentially, rather than being active in only one coding epoch; in fact, transitions between response epochs could be observed in single trials using ensemble statistics. These results suggest that the processing of quality and palatability is likely not performed by independent neural ensembles.

Our study adds to a developing foundation for comparative analysis of mice and rat GC taste processing. Although the anatomies of the two species’ taste systems are similar, little work has been done comparing taste coding between the two species. In all of our current results, we observed a striking similarity between mouse cortical taste coding and that extensively described in rat data (Accolla et al., 2007; Bahar et al., 2004; Fontanini and Katz., 2006; Jones et al., 2007; Piette et al., 2012; Sadacca et al., 2012; Samuelsen et al., 2013; Yamamoto et al., 1985;). Both species code tastes broadly, and these codes reflect identity and taste palatability in separate, sequential epochs. It may well be that differences will emerge with regard to the impact of taste learning on neural coding (which we will examine in the future).

Note that these data do not address the question of whether the coding of taste is necessarily “temporal coding” *per se* (although see Di Lorenzo et al., 2009; Erickson et al., 1994; Hallock and Di Lorenzo, 2006; Lemon and Smith, 2006): mouse GC taste responses become taste-specific at approximately 0.15-0.2 sec after taste delivery, a latency that accords well with the absolute minimum behavioral reaction times to tastes (Halpern and Tapper, 1968, 1971; Perez et al., 2013); firing in that ~0.2-~1.0 epoch is best described as “across-fiber patterns” (although not as “labeled lines,” see Erickson, 1963, 1982). Rather, our data suggest that the term “taste coding” far under-describes the work in which GC is involved, in that GC responses chart the transformation of taste information into an action (i.e., palatability) code. Evidence supporting this conception comes from rat studies showing that the onset of palatability-related firing both predicts (Sadacca et al., 2016) and drives (Li et al., 2016) the onset of taste behavior.

Note that these last pieces of data—the fact that the onset of palatability-related firing precedes and predicts the onset of palatability-related orofacial behaviors (gapes and lateral tongue protrusions)—suggests that this activity is not simply a reflection of said behaviors. And while it might be argued that these oft-studied behaviors might be preceded by other palatability-related behaviors, and that palatability-related firing might be epiphenomenal, efference copy of those motor behaviors, this possibility is vanishingly unlikely for the following reasons: 1) extensive study of taste-related behavior has revealed no palatability-specificity in mouth movements prior to gaping and lateral tongue protrusions (see, e.g., Travers & Norgren, 1986); 2) even if some subtle behaviors have escaped the notice of researchers, it would be surprising if such difficult-to-observe motor acts were to drive such obvious neural activity; and 3) the transition to palatability-related firing does not just precede and predict motor behavior, it is necessary for this behavior to emerge normally (Mukherjee et al., in press). It is much more likely that in the mouse, as in the rat, GC participates not only in “coding” a taste per se, but in reaching decisions regarding how to handle that taste.

Finally, we extend the above-described analyses into the spatial domain (taking this work beyond that addressed in electrophysiological work on rat GC), probing for possible differences in how dorsal/ventral and anterior/posterior sub regions of GC code tastes, because: 1) GC subregions form distinct connections with other brain regions along the dorsal/ventral and anterior/posterior axes (Allen et al., 1991; Haley, et al., 2016; Maffei et al., 2012); and 2) the literature on imaging of mouse GC fails to support consistent conclusions on the matter (Chen et al., 2011; Livneh et al., 2016; Fletcher et al., 2017; Lavi et al., 2018). Our comparisons revealed remarkably little spatial variance of coding, consistent with Fletcher et al (2017; see also Accolla et al., 2007 and Lavi et al., 2018)— neurons across the heart of GC produced similarly broad, temporally complex taste responses.

This invariance in the face of non-uniform input—most notably, dorsal GC receives mainly thalamic input (Allen et al., 1991; Chen et al., 2011; Fletcher et al., 2017), whereas ventral GC receives mainly amygdala input (Allen et al., 1991; Matyas et al., 2014; see Figure 1)—is striking given the difference in information carried in those two pathways; thalamus appears to deliver mainly quality-related firing to GC (Liu et al., 2015; Samuelsen et al., 2013), whereas amygdala-cortical axons are more important for palatability coding (Piette et al., 2012). We have no specific explanation for this lack of difference, although it suggests that GC taste responses are to a large extent a function of distributed network processing, rather than of direct sensory or effective input—i.e., whatever information is carried in each pathway, intra-cortical connectivity ensures that this information is spread across a broader spatial region. This interpretation is supported by the very finding that GC neurons code multiple aspects of tastes in a temporally distributed fashion, which suggests convergence of pathways, and population coding analysis that reveal the temporal codes to be a reliable sequence of coherent network states (see Grossman et al., 2008; Jones et al., 2007; Moran & Katz, 2014; Sadacca et al., 2016). It is also notable, however, that amygdalar inactivation eliminates palatability-related firing in only a majority of GC neurons, enhancing them in the remainder (Piette et al., 2012); perhaps a mapping of this effect would reveal that the palatability-related firing that survives amygdalar inactivation will be localized to dorsal, thalamo-recipient regions of GC.

Together our results shed a new light on the structure of taste coding in mice. They show that GC neurons are broadly tuned, but go on to show that GC neurons code not only multiple tastes but also multiple aspects of taste processing (taste identity and palatability)—the latter in meaningful response dynamics. We also show that these responses are distributed through a broad spatial extent of GC; cortical network processing is distributed, rather than being separated out into spatial subregions. At the largest “big picture” level, these results suggest that the functional unit of sensory processing in GC is ensembles composed of neurons that are responsive to multiple tastes, in which properties of the processing emerge transiently in specific time epochs.

## Acknowledgements

This research was supported by the National Institute on Deafness and Other Communication Disorders (NIDCD) DC006666.

